# Schistosomiasis among pregnant women in the Njombe-Penja health district, Littoral region of Cameroon: A cross sectional study

**DOI:** 10.1101/661900

**Authors:** Calvin Tonga, Charlie Ngo Bayoi, Flore Chanceline Tchanga, Jacqueline Félicité Yengue, Godlove Bunda Wepnje, Hervé Nyabeyeu Nyabeyeu, Lafortune Kangam, Larissa Nono Kouodjip, Patrick Ntonga Akono, Léopold Gustave Lehman

## Abstract

**Background:** Schistosomiasis is a Neglected Tropical Disease with endemic foci in Cameroon. Epidemiological data on schistosomiasis in pregnancy are scarce in the country. This study is about schistosomiasis among pregnant women in the Njombe-Penja health district, where schistosomiasis was reported since 1969.

**Methodology:** Overall, 282 pregnant women were enrolled upon informed consent at first antenatal consultation. A questionnaire was administered to document socio-economic and obstetric information. Stool and terminal urine samples were collected and analysed using the Kato-Katz/formol-ether concentration techniques and centrifugation method respectively. Haemoglobin concentration was measured with finger prick blood, using a URIT-12^®^ electronic haemoglobinometer.

**Principal findings:** The overall prevalence of schistosomiasis was 31.91%. *Schistosoma guineensis, S. haematobium* and *S. mansoni* infections were found in 0.35%, 04.96% and 28.01% of participants respectively. Co-infection with 2 species of *Schistosoma* was found in 04.44% of these women. The prevalence of schistosomiasis was significantly higher in younger women (≤20) and among residents of Njombe. All *S. haematobium* infected women were anemic and infection was associated with significantly lower haemoglobin levels (p=0.02).

**Conclusion:** The prevalence of schistosomiasis is high in pregnant women of the Njombe-Penja health district, with possible adverse pregnancy outcomes. Female of childbearing age should be considered for mass drug administration.

**Author summary:** Pregnant women are known to be more vulnerable to infectious diseases and in their case, at least two lives are at risk. Although schistosomiasis remains a major public health issue in Cameroon, epidemiological data on schistosomiasis in pregnancy are scarce. These data are of high interest for informed decision-making. We examined stools and urines from 282 women of the Njombe-Penja Health district and measured their blood levels. Overall, 31.91% of women were infected, mostly younger ones and those living in the town of Njombe. Three species of Schistosoma parasite were identified. Women having urinary schistosomiasis had lower blood levels. These results show that the prevalence of schistosomiasis is high in pregnant women of Njombe. Also, because of the anemia it induces, the disease can lead to adverse pregnancy outcomes on the woman and her foetus. Treating female of childbearing age would cure the disease and prevent adverse outcomes.

## Introduction

Schistosomiasis also known as bilharzia is an acute and chronic neglected tropical disease (NTD) caused by parasitic flatworms of the genus *Schistosoma*. Scistosomiasis affects at least 240 million individuals accross 74 countries and territories worldwide, including 40 million women of child bearing age, mostly the underprivileged [1–2]. Sub-Saharan Africa accounts for about 70% of cases, most regions and islands being affected, except for Kalahari desert and the extreme southern part of the continent [3–4]. Six species affect man, namely *Schistosoma haematobium* that lives within the perivesicular venules, *Schistosoma mansoni, Schistosoma guineensis, Schistosoma intercalatum, Schistosoma japonicum* and *Schsistosoma mekongi*, that live within the mesenteric venules. In Africa, schistosomiasis is mostly caused by *S. haematobium* and *S. mansoni* [5,6].

In a survey carried-out in 1999, it was estimated that 1 066 000 persons were infected with *S. haematobium* with a national prevalence of 6.1%; 713 000 persons were affected by *Schistosoma mansoni* with a national prevalence of 4.5%. About ten thousands new cases are recorded each year [7–9]. Endemic foci of *S. haematobium, S. guineensis* and *S. mansoni* are mostly found in the northern part of the country and the South-West region. In addition, there are spotted foci including that in the Moungo Division of the Littoral region [9–12].

Tremendous efforts have been made since 2004 to fight against schistosomiasis and other NTDs in Cameroon. National and international commitment is perceptible through activities of dedicated research institutions and the control programme. Since 2007, national integrated deworming campaigns are implemented yearly and praziquantel is distributed in endemic areas [9,12]. However, mass drug administration (MDA) of praziquantel against schistosomiasis targets only children of 1-14 years old; adults including pregnant women are generally not considered for treatment, though there are evidence of increased vulnerability to the infection during pregnancy. Moreover, the availability of Praziquantel within the health system remains poor [13–15].

Epidemiological data on schistosomiasis during pregnancy are scarce in sub-Saharan Africa as most of studies focus on schoolchildren [16]. In Cameroon as in other sub-Saharan African countries, few studies on maternal schistosomiasis have been reported [17–20]. It is thus difficult to have a reliable estimate of the extent of the infection in this stratum of the population. Moreover, being left untreated during control interventions, pregnant women could face adverse pregnancy outcome and serve as reservoirs of transmission, thus jeopardizing control efforts over time.

The present paper reports on Schistosomiasis among pregnant women in the Njombe-Penja health district, where schistosomiasis was described since 1969 [21,22]. It presents the prevalence, factors associated with the disease and effects on blood levels in pregnant women.

## Methods

### Ethical considerations

Ethical clearance was obtained from the University of Douala Institutional Review Board (CEI-DU/153/02/2015/T) and an administrative clearance was issued by the Regional Delegation of Public Health for the Littoral region (593/L/MINSANTE/DRSPL/BCASS). Pregnant women were approached during first contact with the health facility for pregnancy follow-up. The objectives of the study were explained to them in the language they understood and their questions were answered. Only volunteer pregnant women who signed the informed consent form were enrolled. Participants found infected were referred to clinicians of the health centre for adequate care. There was no difference in the care provided to pregnant women who accepted to participate in the study and those who did not.

### Study area and population

This cross sectional study was conducted from April to December 2016 in Njombe (Njombe-Penja Health District, Moungo Division, Littoral region of Cameroon), a semi-urban setting situated at 94Km west of Douala the economic capital city of Cameroon. The coordinates of Njombe ranged from altitude 99 metres, latitude 4°34’50” N and longitude 9°39’52” E. The study area is subject to a tropical climate type with a long rainy season and a short dry season. The average temperature is 26.6 °C and averages rainfall is 3002 mm. In 2016, total population of the Njombe-Penja health district was estimated at 47 839 inhabitants, with about 1533 pregnant women [23]. This is a farming area, with food crops and fruits farms adjacent to major industrial banana, white pepper and flowers farms.

The study population consisted of volunteer pregnant women visiting the Njombe 1 Integrated Health Centre, who signed inform consent forms. In this area, Ngolle reported a prevalence of 24% for urinary schistosomiasis in senior primary school children while Lehman *et al*. reported a prevalence of of 13% for intestinal schistosomiasis in the general population [24,25]. The highest prevalence was used for sample size calculation using the Lorentz formula as follows: 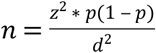 and the minimum sample size was estimated at 280. Overall, 282 pregnant women were enrolled in this study.

### Study design, information and sample collection

This study was a cross-sectional survey carried out from April to December 2016. Pregnant women received during first antenatal consultation (ANC) were enrolled, prior to any health intervention related to their pregnancy. A structured questionnaire was administered in the language they understood best (French, English or Pidgin), in order to collect socio-economic and obstetric information. Each woman received a labelled stool container for stool sample collection and a labelled graduated plastic urine container in which she provided 30 to 50 mL of urine. All urine samples were collected between 10 a.m. and 2 p.m. to coincide with the peack of excretion of *S. haematobium* eggs. Peripheral blood sample for determination of haemoglobin concentration was collected from finger prick.

### Laboratory analysis

Analysis of capillary blood was done immediately with a URIT-12^®^ electronic haemoglobinometer, for measurement of haemoglobin. Women with haemoglobin levels lower than 11g/dL were considered anaemic [26].

The urine samples were transported to the laboratory and analysed using the centrifugation method as described by Okanla [27]. Briefly, each urine container was shaken and 10 mL of urine were drawn and transferred into a 20 mL centrifuge tube. The tubes were centrifuged at 1000 rpm for 5 minutes. The supernatant was discarded and one drop of the sediment was put on a clean glass slide; a drop of Lugol’s iodine was added. The preparation was then cover-slipped and examined under the x20 and x40 objective lenses of a CyScope^®^ light microscope. Urinary schistosomiasis status was determined by the detection of characteristic *S. haematobium* eggs in urine. Intensity of infection (egg/10mL) was estimated by multiplying the number of eggs counted in one drop times 20. Mild infection was defined as less than 50 ova per 10 mL urine and severe infection as more than 50 ova per 10 mL urine [28].

Kato-Katz (KK) and Formol-Ether (FE) techniques were used for stool samples analyses and viewed at x20 and x40 magnification as described by WHO [29]. For the KK technique, each sample was sieved and calibrated using the KK template in order to obtain about 41.7 mg of sieved stool on the slide that was then covered with a glycerol-malachite green impregnated cellophane strip. The preparation was examined 24 to 48 hours later for schistosome eggs. For the FE technique, stool was weighted on an electronic scale and 1 gr was mixed in 10 mL of 10% buffered formaline; after filtration with gauze, 7 mL of the preparation was mixed with 3 mL of ether and the mixture was centrifuged at 1000 rpm for 5 minutes. The supernatant was then discarded and a sediment of about 500μL left at the bottom of the tube. One drop of the sediment was mixed with a drop of Lugol’s iodine on a clean glass slide. The preparation was then cover-slipped and examined. The number of eggs for each parasite was counted; the intensity of infection, expressed as eggs per gram of faeces (epg), was calculated by multiplying the egg count by 24 for KK and 10 for FE. Mild infection was defined as <100 epg, moderate infection as 101-400 epg and severe infection as >400 epg) [30].

### Statistical analysis

From the information collected through the questionnaires, participants were categorized according to their age groups (≤20 years and >20 years), level of education (≤primary school, ≥secondary school), marital status (single, married), pregnancy age (≤28 weeks and >28 weeks), monthly income (based on the Interprofessional guaranteed minimum wage or SMIG which is 36 270 FCFA; <72540 and ≥72540). According to number of pregnancies (gravidity), they were categorized as paucigravida (less than 4 pregnancies) and multigravida (4 pregnancies and more). All data were entered in an Excel worksheet and analysed using Epi Info version 7.2.1.0., 2017, CDC. Geometric egg density was calculated for each of the parasite after log transformation of laboratory results. The sensitivity of laboratory techniques used for the detection of intestinal schistosomiasis was calculated through dividing the number of positive cases obtained with this specific technique by the total of positive cases from both techniques. Bivariate and multivariate analyses were performed to assess association between exposure and outcome variables. In bivariate analysis, Chi-square and Fisher’s exact probability tests were used to assess differences in proportions; ANOVA was used to assess differences in means and Yates Odds Ratios (OR) were calculated to compare the susceptibility of individuals or groups to different parameters. Backward stepwise binary logistic regression was used for multivariate analysis, including variables with p-value ≤0,5 at bivariate analysis [31]. Missing data were not computed. The level of significance was set at p=0.05.

## Results

### Description of study participants

Overall, 282 pregnant women participated into this study, mostly residents of the town of Njombe (90.07%) with mean age of 24.8±5.6 (range: 14–43) years. The mean haemoglobin concentration was 10.0±1.2 (range: 7.2–13.9) g/dL and mean number of pregnancies was 3±2 (range: 1–9). Among the participants, 5 (1.77%) were infected with HIV (Table 1). Most of the participants (48.53%) were farmers or housewives and 87.62% had less than twice the SMIG (Interprofessional guaranteed minimum wage) as monthly revenue for their families.

**Table 1.**
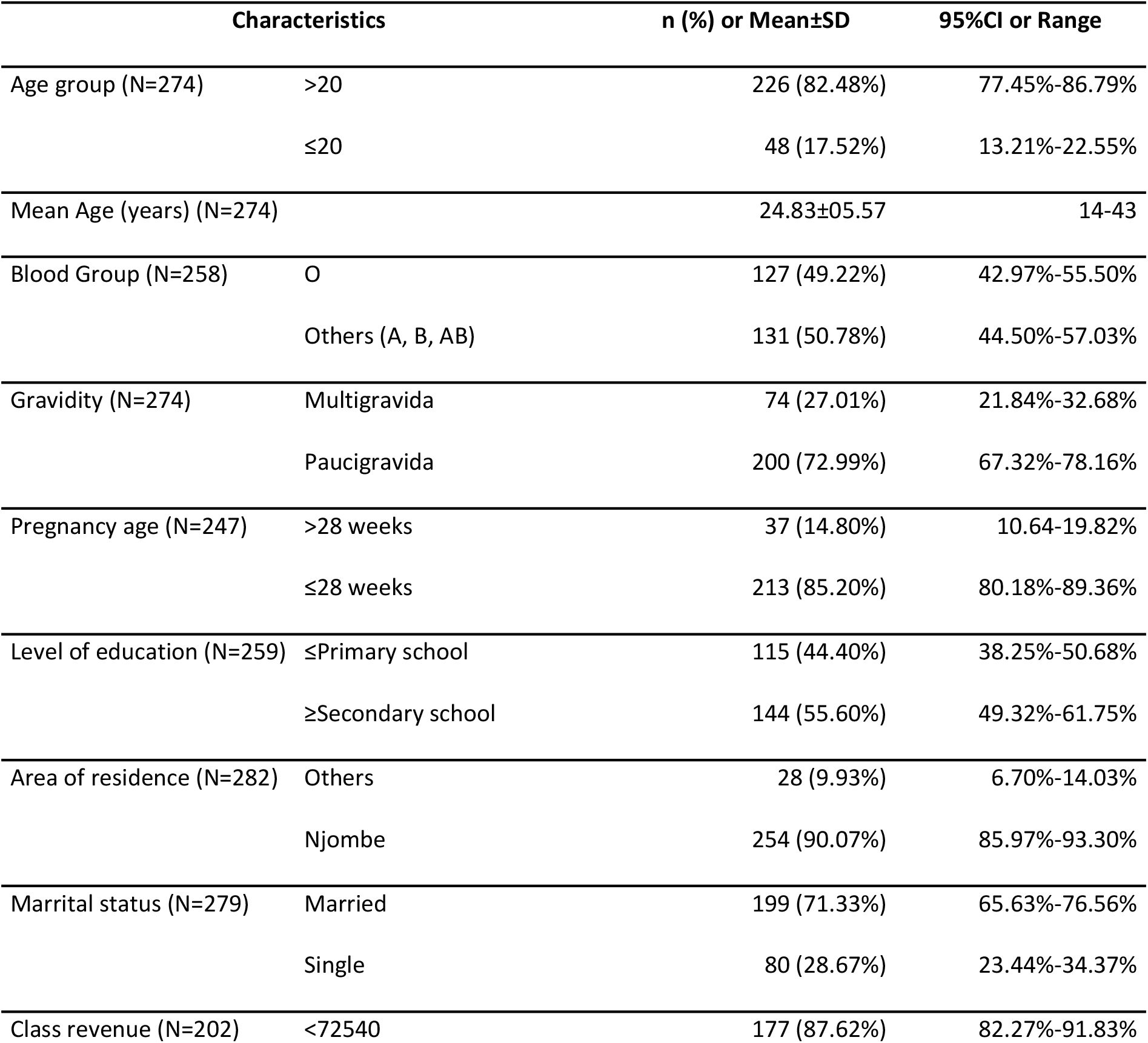
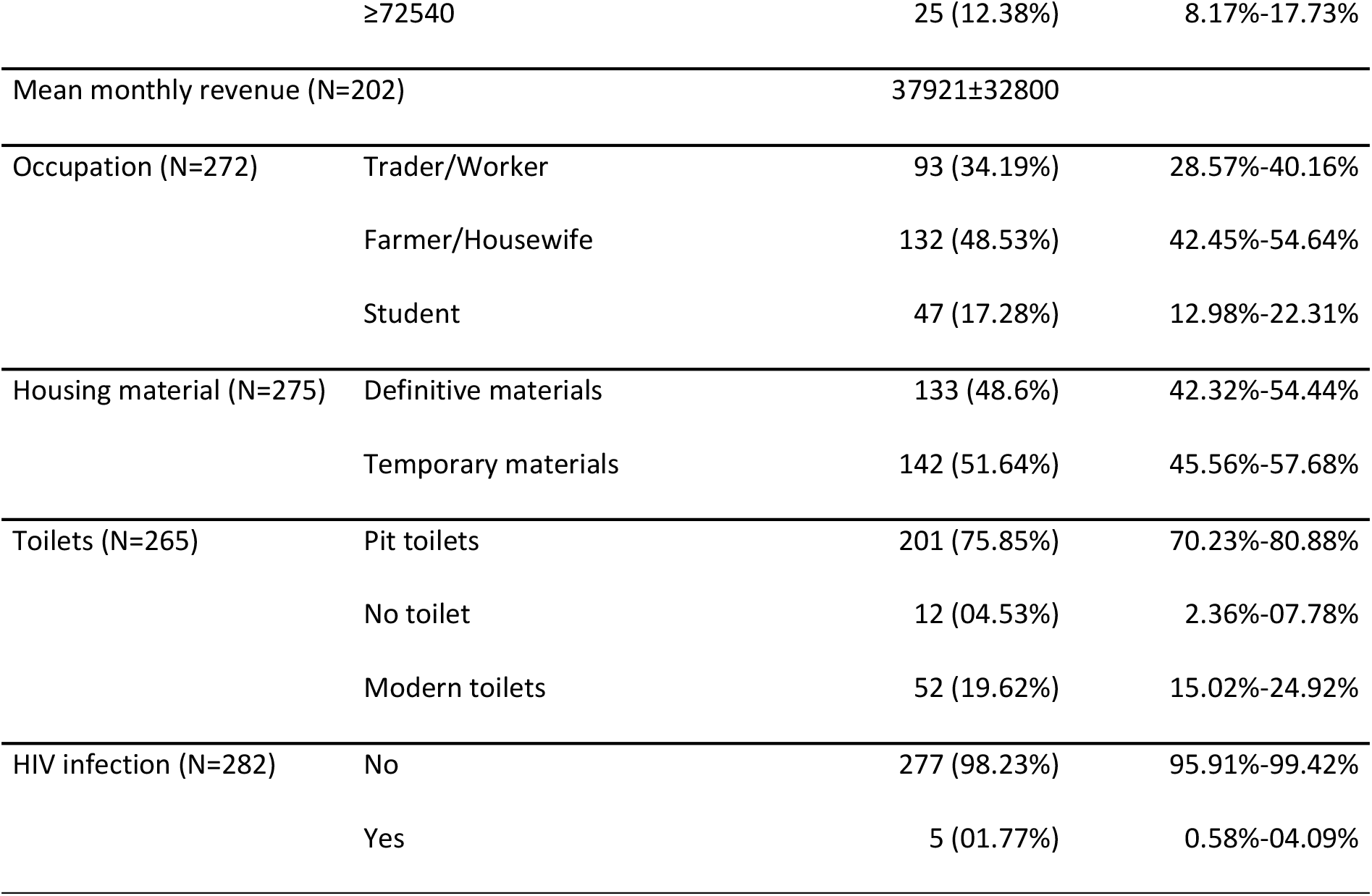
Characteristics of the study population

| Characteristics | | n (%) or Mean±SD | 95%CI or Range |
| --- | --- | --- | --- |
| Age group (N=274) | >20 | 226 (82.48%) | 77.45%-86.79% |
| | $\leq 20$ | 48 (17.52%) | 13.21%-22.55% |
| Mean Age (years) (N=274) | | 24.83±05.57 | 14-43 |
| Blood Group (N=258) | O | 127 (49.22%) | 42.97%-55.50% |
|  | Others (A, B, AB) | 131 (50.78%) | 44.50%-57.03% |
| Gravidity (N=274) | Multigravida | 74 (27.01%) | 21.84%-32.68% |
|  | Paucigravida | 200 (72.99%) | 67.32%-78.16% |
| Pregnancy age (N=247) | >28 weeks | 37 (14.80%) | 10.64-19.82% |
| | $\leq 28$ weeks | 213 (85.20%) | 80.18%-89.36% |
| Level of education (N=259) | ≤Primary school | 115 (44.40%) | 38.25%-50.68% |
| | ≥Secondary school | 144 (55.60%) | 49.32%-61.75% |
| Area of residence (N=282) | Others | 28 (9.93%) | 6.70%-14.03% |
|  | Njombe | 254 (90.07%) | 85.97%-93.30% |
| Marrital status (N=279) | Married | 199 (71.33%) | 65.63%-76.56% |
|  | Single | 80 (28.67%) | 23.44%-34.37% |
| Class revenue (N=202) | <72540 | 177 (87.62%) | 82.27%-91.83% |
|  | ≥72540 | 25 (12.38%) | 8.17%-17.73% |
| Mean monthly revenue (N=202) |  | 37921±32800 |  |
| Occupation (N=272) | Trader/Worker | 93 (34.19%) | 28.57%-40.16% |
|  | Farmer/Housewife | 132 (48.53%) | 42.45%-54.64% |
|  | Student | 47 (17.28%) | 12.98%-22.31% |
| Housing material (N=275) | Definitive materials | 133 (48.6%) | 42.32%-54.44% |
|  | Temporary materials | 142 (51.64%) | 45.56%-57.68% |
| Toilets (N=265) | Pit toilets | 201 (75.85%) | 70.23%-80.88% |
|  | No toilet | 12 (04.53%) | 2.36%-07.78% |
|  | Modern toilets | 52 (19.62%) | 15.02%-24.92% |
| HIV infection (N=282) | No | 277 (98.23%) | 95.91%-99.42% |
|  | Yes | 5 (01.77%) | 0.58%-04.09% |

### Diversity, prevalence, intensity of Schistosoma infection

Three species of the *Schistosoma* genus were found in this study, namely *S. guineensis, S. haematobium* and *S. mansoni. S. guineensis* was detected only with the FE technique but not with the KK technique. Overall, 15 cases of all *S.mansoni* infections were missed with the KK technique. However, the KK technique was more sensitive for the detection of *S. mansoni* than the Formol-Ether technique; the number of *S. mansoni* infections detected using KK was about double of that detected with FE (KK sensitivity: 81.01%; FE sensitivity: 45.57%). Moreover, egg count with KK was overall higher than with FE (GMED: 55 epg versus 52 epg), although not statistically significant (H=12.82, p=0.17; Mann-Whitney test; table 2).

**Table 2.**
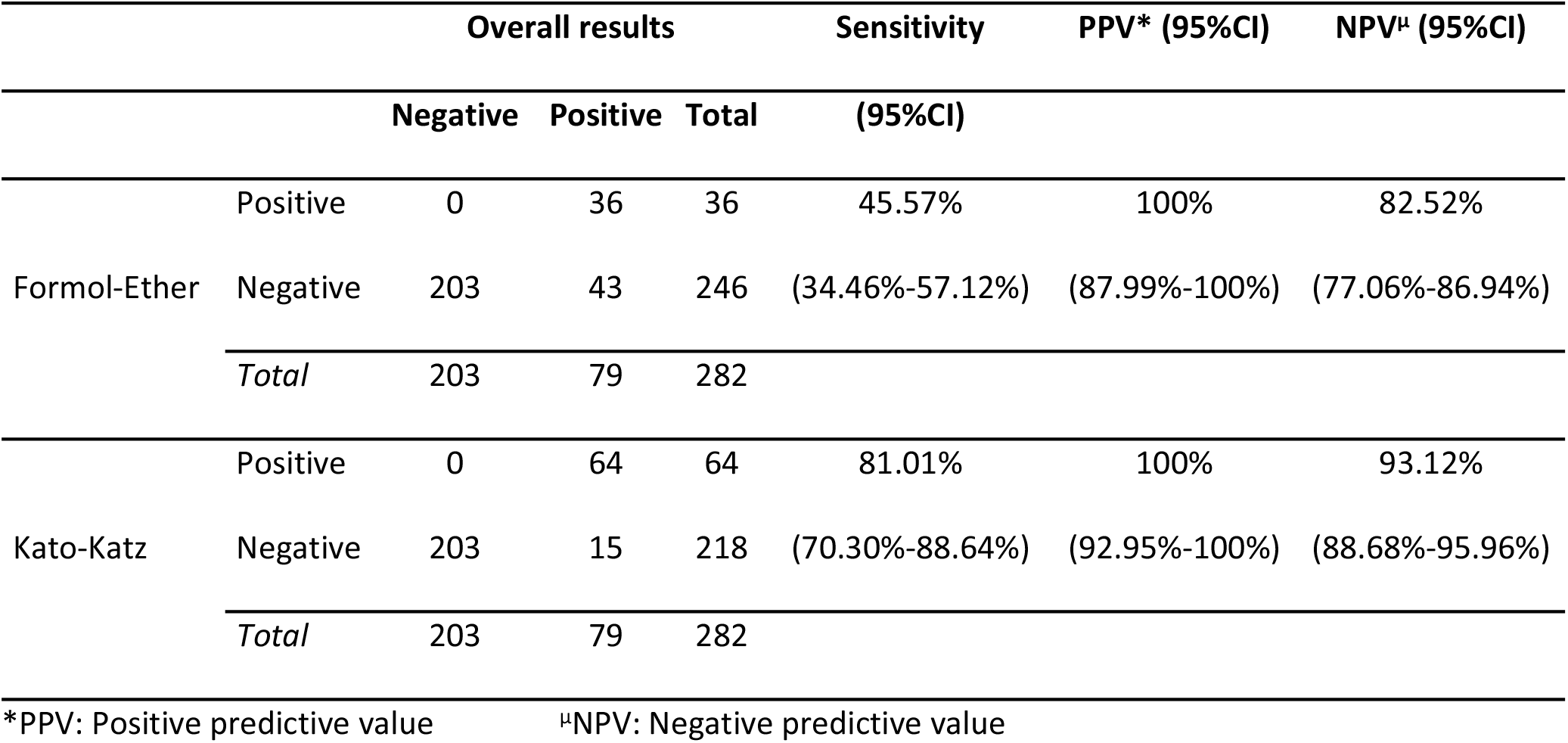
Sensitivity of Formol-Ether and Kato-Katz techniques for the detection of *S. mansoni* infections

Ninety women (31.91%) were found infected with *Schistosoma. S. mansoni* was the most prevalent species (28.01%), followed by *S. haematobium* (4.96%). Only one case of *S. guineensis* infection was detected, with the egg count corresponding to mild infection. For *S. haematobium*, 50% of infections were severe whereas for *S. mansoni*, severe, moderate and mild infections were 5.06%, 18.99% and 75.95% respectively (table 3).

**Table 3.** Prevalence of Schistosoma species and geometric mean egg densities

| Species | Technique | Frequency | GMED* |
| --- | --- | --- | --- |
| *Schistosoma guineensis* | Formol-Ether | 1 (0.35%) | 25±0 |
|  | Kato-Katz | 0 (0.0%) | 0 |
|  | *Total* | **1 (0.35%)** | **25±0** |
| *Schistosoma mansoni* | Formol-Ether | 36 (12.77%) | 52±2 |
|  | Kato-Katz | 64 (22.70%) | 55±2 |
|  | *Total* | **79 (28.01%)** | **54±2** |
| *Schistosoma haematobium* | CM<sup>u</sup> | **14 (4.96%)** | **51±2** |
| *Schistosoma spp.* |  | **90 (31.91%)** |  |

<sup>u</sup>CM: Centrifugation Method \*GMED: Geometric Mean Egg density (egg per gram and egg per 10mL)

Four cases of bi-infection were found and the sole *S. guineensis* detected was in a woman also infected with *S. mansoni* (table 4).

**Table 4.** Frequency of mono and bi-infections with Schistosoma spp.

| Type of infection |  | Number of cases | Frequency (95%CI) |
| --- | --- | --- | --- |
| Mono-infection | *S. haematobium* | 11 | 12.22% (06.96%-20.57%) |
|  | *S. mansoni* | 75 | 83.33% (74.31%-89.63%) |
|  | Total | 86 | 95.56% (89.12%-98.26%) |
| Bi-infection | *S. guineensis_S. mansoni* | 1 | 1.11% (02.20%-06.03%) |
|  | *S. haematobium_S. mansoni* | 3 | 3.33% (01.14%-09.35%) |
|  | Total | 4 | 4.44% (01.74%-10.88%) |

### Factors associated with schistosomiasis

As shown in table 5, bivariate analysis revealed a significant association of blood group, level of education and occupation with *S. haematobium* infection. The prevalence of *S. haematobium* infection was about 4 folds higher in women of group O compared with other blood groups, and 5 folds higher in more educated women. Students were more affected than other occupational categories. In addition, *S. haematobium* was found exclusively in women living in Njombe. However, none of these factors was significantly associated with *S. haematobium* infection in logistic regression.

**Table 5.** Factors associated with Schistosoma infection in the study population

| Characteristic | Class | *Schistosoma haematobium* |  |  | *Schistosoma mansoni* |  |  | All schistoma species |  |  |
| --- | --- | --- | --- | --- | --- | --- | --- | --- | --- | --- |
|  |  | Infected n(%) | OR (CI 95%) | p-value* | Infected n(%) | OR (CI 95%) | p-value | Infected n(%) | OR (CI 95%) | p-value |
| Age group | >20 | 12 (5.31%) | 1.29 (0.28-5.96) | 0.99 | 57 (25.22%) | 2.12 (1.11-4.05) | **0.034** | 67 (29.65%) | 1.85 (0.98-3.49) | 0.08 |
|  | ≤20 | 2 (4.17%) |  |  | 20 (41.67%) |  |  | 21 (43.75%) |  |  |
| Blood Group | O | 11 (8.40%) | 3.79 (1.02-13.92) | **0.032** | 41 (31.30%) | 1.25 (0.73-2.14) | **0.51** | 49 (37.40%) | 1.45 (0.86-2.45) | 0.20 |
|  | Others (A, B, AB) | 3 (2.36%) |  |  | 34 (26.77%) |  |  | 37 (29.13%) |  |  |
| Gravidity | Multigravida | 2 (2.70%) | 2.29 (0.56-15.42) | 0.43 | 16 (21.62%) | 1.55 (0.83-2.92) | 0.22 | 18 (24.32%) | 1.64 (0.90-3.00) | 0.14 |
|  | Paucigravida | 12 (6.00%) |  |  | 60 (30.00%) |  |  | 69 (34.50%) |  |  |
| Pregnancy age | >28 weeks | 3 (8.11%) | 0.50 (0.13-2.40) | 0.55 | 12 (32.43%) | 0.76 (0.36-1.62) | 0.61 | 14 (37.84%) | 0.71 (0.34-1.46) | 0.45 |
|  | ≤28 weeks | 9 (4.23%) |  |  | 57 (26.76%) |  |  | 64 (30.05%) |  |  |
| Level of education | ≤Primary school | 2 (1.74%) | 4.67 (1.01-21.52) | **0.031** | 30 (26.09%) | 1.05 (0.60-1.83) | 0.97 | 32 (27.83%) | 1.30 (0.76-2.21) | 0.41 |
|  | ≥Secondary school | 11 (7.64%) |  |  | 39 (27.08%) |  |  | 48 (33.33%) |  |  |
| Area of residence | Others | 0 (0%) | NA | 0.37 | 2 (7.14%) | 5.66 (1.31-24.42) | **0.02** | 2 (07.14%) | 6.89 (1.60-29.71) | **0.006** |
|  | Njombe | 14 (5.51%) |  |  | 77 (30.31%) |  |  | 88 (34.65%) |  |  |
| Marrital status | Married | 9 (4.52%) | 1.41 (0.41-4.34) | 0.55 | 50 (25.13%) | 1.52 (0.86-2.67) | 0.19 | 56 (28.14%) | 1.70 (0.99-2.93) | 0.07 |
|  | Single | 5 (6.25%) |  |  | 27 (33.75%) |  |  | 32 (40.00%) |  |  |
| Class revenue | <72540 | 7 (3.95%) | 0.47 (0.09-2.42) | 0.60 | 49 (27.68%) | 0.82 (0.31-2.19) | 0.88 | 54 (30.51%) | 1.07 (0.44-2.63) | 1 |
|  | ≥72540 | 2 (8.00%) |  |  | 6 (24.00%) |  |  | 8 (32.00%) |  |  |
| Occupation | Trader/Worker | 3 (3.23%) | 0.19 (0.05-0.77) | **0.017** | 31 (33.33%) | 1.31 (0.50-2.83) | 0.62 | 32 (34.41%) | 0.77 (0.38-1.59) | 0.61 |
|  | Farmer/Housewife | 4 (3.03%) | 0.18 (0.05-0.64) | **0.008** | 31 (23.48%) | 0.80 (0.38-1.71) | 0.71 | 35 (26.52%) | 0.53 (0.26-1.07) | 0.11 |
|  | Student | 7 (14.89%) | NA |  | 13 (27.66%) |  |  | 19 (40.43%) |  |  |
| Housing material | Definitive materials | 5 (3.76%) | 1.73 (0.56-05.83) | 0.49 | 36 (27.07%) | 1.09 (0.64-1.85) | 0.84 | 40 (30.08%) | 1.19 (0.71-1.97) | 0.59 |
|  | Temporary materials | 9 (6.34%) |  |  | 41 (28.87%) |  |  | 48 (33.80%) |  |  |
| Toilets | Pit toilets | 10 (4.98%) | 0.86 (0.23-3.23) | 0.99 | 56 (27.86%) | 0.95 (0.49-1.87) | 1 | 63 (31.34%) | 0.86 (0.45-1.64) | 0.78 |
|  | No toilet | 1 (8.33%) | 1.48 (0.14-15.66) | 0.99 | 4 (33.33%) | 1.23 (0.32-4.72) | 1<sup>f</sup> | 5 (41.67%) | 1.35 (0.37-4.86) | 0.74<sup>f</sup> |
|  | Modern toilets | 3 (5.77%) | NA |  | 15 (28.85%) |  |  | 18 (34.62%) |  |  |
| HIV infection | No | 13 (4.69%) | 0.20 (0.02-1.89) | 0.23 | 78 (28.16%) | 0.64 (0.07-5.80) | 1 | 88 (31.77%) | 1.43 (0.23-8.72) | 1<sup>f</sup> |
|  | Yes | 1 (20.00%) |  |  | 1 (20.00%) |  |  | 2 (40.00%) |  |  |
| OVERALL |  | 14 (4.96%) |  |  | 79 (17.94%) |  |  | 90 (31.91%) |  |  |

\*Except when specified, all p-values are calculated from corrected (Yates) Chi-2.
<sup>f</sup>: p-value from Fisher exact test

Infection with *S. mansoni* was significantly associated with age group and residence area. Women aged 20 or less were twice as infected as their older counterparts; mean age of infected women was significantly lower than that of their non-infected counterparts (23.7 years versus 25.3 years; p=0.04). The prevalence of the infection in women living in Njombe was 6 folds higher than that in women living in other settings. This was later confirmed in logistic regression (aOR=2.06, p=0.03; aOR=5, p=0.03 respectively).

Considering all Schistosoma species, infection was significantly associated with area of residence, women living in Njombe being more infected (OR=6.89; p=0.006). Also, mean age was significantly low for infected women (23.8 years versus 25.3 years; p=0.029). This result was confirmed with logistic regression (aOR=5.96, p=0.02).

### Influence of schistosomiasis on blood levels

The prevalence of anaemia in women infected with *S. haematobium* was 100%. Moreso, mean haemoglobin concentration was lower in infected women (9.26 versus 10.08; p=0.02). The prevalence of anaemia as well as haemoglobin levels were not significantly affected by *S. mansoni* infection (table 6).

**Table 6.**
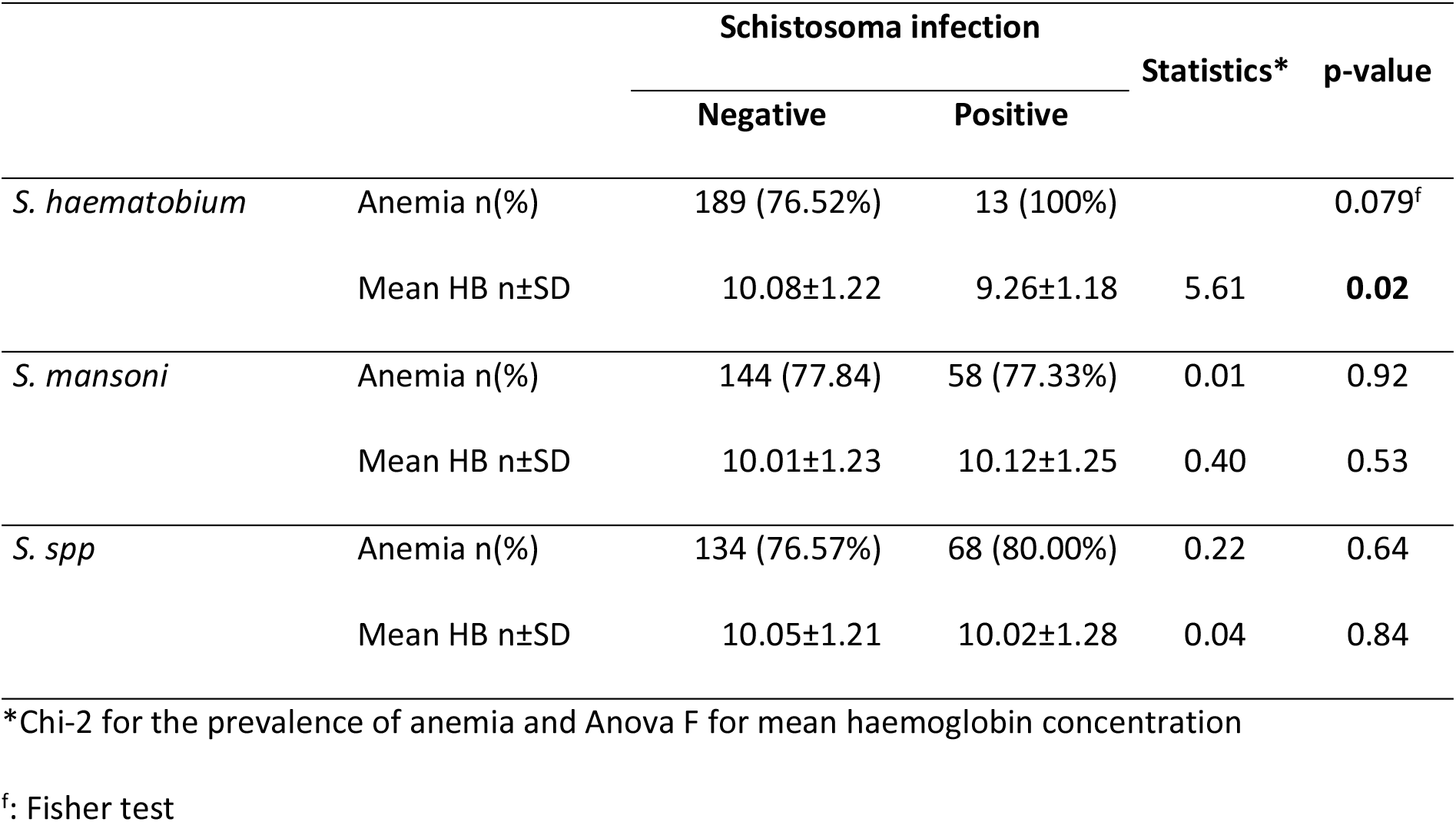
Effect of schistosomiasis on the prevalence of anemia and haemoglobin concentration

|  |  | Schistosoma infection |  | Statistics* | p-value |
| --- | --- | --- | --- | --- | --- |
|  |  | Negative | Positive |  |  |
| *S. haematobium* | Anemia n(%) | 189 (76.52%) | 13 (100%) |  | 0.079<sup>f</sup> |
|  | Mean HB n±SD | 10.08±1.22 | 9.26±1.18 | 5.61 | **0.02** |
| *S. mansoni* | Anemia n(%) | 144 (77.84) | 58 (77.33%) | 0.01 | 0.92 |
|  | Mean HB n±SD | 10.01±1.23 | 10.12±1.25 | 0.40 | 0.53 |
| *S. spp* | Anemia n(%) | 134 (76.57%) | 68 (80.00%) | 0.22 | 0.64 |
|  | Mean HB n±SD | 10.05±1.21 | 10.02±1.28 | 0.04 | 0.84 |

\*Chi-2 for the prevalence of anemia and Anova F for mean haemoglobin concentration
<sup>f</sup>: Fisher test

## Discussion

This study aimed at determining the prevalence and diversity of schistosomiasis among pregnant women in the Njombe-Penja health district, potential risk factors for the disease as well as its influence on blood levels. It brings additional epidemiological data on schistosomiasis in adults and especially pregnant women, known to be more vulnerable to most infectious diseases with at least two lives at risk.

Three species of Schistosoma were found in our study population; they had been described in the country by Ratard *et al*. [32]. The overall prevalence of Schistosoma infection was 31.91%; thus, about 1/3 of pregnant women are infected. *S. guineensis* prevalence was very low, marking the gradual disappearance of this species in the Njombe area because of interspecific competition and introgressive hybridization leading to its replacement by *S. haematobium* [8,33,34].

The prevalence of urinary schistosomiasis among pregnant women was 4.96%, not far from the national estimate of 6.1% [7]. Our result is similar to that reported by Siegrist and Siegrist-Obimpeh in Ghana (4.5%) [35]. It is however lower than that reported in Nigeria by Eyo *et al*., Salawu and Odaibo as well as the 46.8% found in Munyengue, South-West of Cameroon by Anchang-Kimbi *et al*. [16,18,20]. The study population in Munyengue (Cameroon) and in Nigeria were highly exposed to *S. haematobium* infection, due to their absolute dependence on natural water sources for domestic activities and bathing. In the Njombe-Penja health district, there is a pipe network for the supply of potable water, which to some extent reduces the contact with natural water bodies, thus the exposure to infective worms. In addition, as reported by Tchuem Tchuente *et al*. [12], the number of *S. haematobium* high transmission foci have increased in the South-West region as compared with findings from the 1985-1987 mapping by Ratard *et al*. [32]. In fact, since 2005, Mass Drug Administration (MDA) for schoolchildren with praziquantel was implemented in two districts; Loum health district in the Littoral region (divided into Loum and Njombe-Penja Health districts in 2012) and Mbonge in the South-West region. Munyenge that belongs to the Muyuka health district was thus not concerned. *S. haematobium* transmission have probably increased in the Munyengue area, Muyuka health district while Loum and Njombe-Penja health districts in the Littoral region experienced a decline in the prevalence and transmission of the disease. Another possible explanatory could be difference in the abundance of the *Bulinus* snail intermediate host.

The prevalence of *S. mansoni* was 28.01%, higher than that reported by Lehman *et al*. and Payne *et al*. in this same setting [25,36]. Our study population was aged 14 to 43 years while theirs were 0 to 77 years and 3 to 78 years respectively. The difference could be due to the age range of study population as Mass Drug Administration (MDA) targets primary schoolchildren and our study participants were thus excluded. However, the prevalence of *S. mansoni* was more than five folds higher than that of *S. haematobium*. This might be due to the many factors including lower sensitivity of *S. mansoni* to Praziquantel in participants who were treated during previous campaigns [37), insufficient coverage of MDA in the population in past years [38]. Also, differences in the abundance of host snails as well as in the transmission pattern of both species in the area should be considered.

*S. haematobium* infection was found only in women living in Njombe and the prevalence of *S. mansoni* infection was about 7 folds higher in women living in Njombe than their counterparts from neighbouring settings. This shows that the transmission area for *S. mansoni* is wider than that of *S. haematobium*, and Njombe is the epicentre of schistosomiasis in the Njombe-Penja health district. The prevalence of shistosomiasis was higher in younger women. This is in agreement with previous studies in Cameroon [20]. It is consistent with the general pattern in helminthic infections; the prevalence increases with age from infancy to adolescence, reaches a peak around the age of 19 then starts declining [39]. Older women would more often send their children or younger relatives, for activities involving contact with water bodies. Age dependent immunity to *S. haematobium*, has also been shown to affect the prevalence and egg output in infected persons [40].

Infection with *S. haematobium* had a negative effect, causing a reduction of blood levels. The contribution of schistosomiasis in inducing anaemia in pregnant women has been documented [16]. The findings in this study are consistent with previous studies. Four mechanisms by which shistosome infections lead to anemia have been suggested. They include extracorporeal loss of iron through frank or occult hemorrage due to egg passage across the intestinal or the bladder wall; sequestration and hemolysis of red blood cells in a context of splenomegaly due to portal hypertension; elevated levels of pro-inflammatory cytokines in response to infection with subsequent drop in erythropoiesis; autoimmune hemolysis of red blood cells [41]. Anemia exposes the pregnant woman and her foetus to adverse pregnancy effects such as maternal death and low-birth weight with subsequent infant mortality.

Limitations of our study are that we did not investigate contact with water bodies as a determinant of schistosomiasis and the association of schistosomiasis with anaemia left out possible infection with other blood parasites.

## Conclusion

Schistosomiasis was found in about one-third of pregnant women. Younger women and women residing in the town of Njombe were more affected. Infected women had lower blood levels, which could lead to adverse pregnancy outcomes. This calls for the continuation and intensification of ongoing control interventions including adequate water supply, sanitization efforts, mass drug administration in schoolchildren and behavioural change for the elimination of the disease in the area. Also, the national control program should consider female of childbearing age for mass drug administration.

## Acknowledgement

We would like to thank all the women who volunteered for participating in this study and all those who contributed in any way to the success of this study. Our sincere gratitude goes to Mrs Basua Rebecca, Chief of the Njombe 1 Integrated Health Centre and the entire staff of this health facility for their collaboration.

## Supporting Information Legends

S1 Checklist: STROBE Checklist

S2 S. mansoni: Picture of *Schistosoma mansoni* egg on a Kato-Katz slide

S3 S. haematobium: Picture of *Schistosoma haematobium* egg on a formol-ether slide

S4 S. haematobium 2: Picture of *Schistosoma haematobium* egg on a formol-ether slide

S5 S. guineensis: Picture of *Schistosoma guineensis egg* on a formol-ether slide

